# The inner membrane protein YhdP modulates the rate of anterograde phospholipid flow in *Escherichia coli*

**DOI:** 10.1101/2020.08.09.213157

**Authors:** Jacqueline Grimm, Handuo Shi, Wei Wang, Angela M. Mitchell, Ned S. Wingreen, Kerwyn Casey Huang, Thomas J. Silhavy

## Abstract

The outer membrane (OM) of Gram-negative bacteria is a selective permeability barrier that allows uptake of nutrients while simultaneously protecting the cell from harmful compounds. The basic pathways and molecular machinery responsible for transporting lipopolysaccharides (LPS), lipoproteins, and β-barrel proteins to the OM have been identified, but very little is known about phospholipid (PL) transport. To identify genes capable of affecting PL transport, we screened for genetic interactions with *mlaA*\*, a mutant in which anterograde PL transport causes the inner membrane (IM) to shrink and eventually rupture; characterization of *mlaA*\*-mediated lysis suggested that PL transport can occur via a high-flux, diffusive flow mechanism. We found that YhdP, an IM protein involved in maintaining the OM permeability barrier, modulates the rate of PL transport during *mlaA*\*-mediated lysis. Deletion of *yhdP* from *mlaA*\* reduced the rate of IM transport to the OM by 50%, slowing shrinkage of the IM and delaying lysis. As a result, the weakened OM of Δ*ydhP* cells was further compromised and ruptured before the IM during *mlaA*\*-mediated death. These findings demonstrate the existence of a high-flux, diffusive pathway for PL flow in *Escherichia coli* that is modulated by YhdP.

**Significance Statement:** The outer membrane (OM) of Gram-negative bacteria serves as a barrier that protects cells from harmful chemical compounds, including many antibiotics. Understanding how bacteria build this barrier is an important step in engineering strategies to circumvent it. A long-standing mystery in the field is how phospholipids (PLs) are transported from the inner membrane (IM) to the OM. We previously discovered that a mutation in the gene *mlaA* causes rapid flow of PLs to the OM, eventually resulting in IM rupture. Here, we found that deletion of the gene *yhdP* delayed cell death in the *mlaA* mutant by slowing flow of PLs to the OM. These findings reveal a high-flux, diffusive pathway for PL transport in Gram-negative bacteria modulated by YhdP.

## Introduction

The outer membrane (OM) of Gram-negative bacteria is an asymmetric bilayer composed of lipopolysaccharides (LPS) in the outer leaflet and PLs in the inner leaflet (1). Strong lateral interactions between LPS molecules in the outer leaflet result in a bilayer that is impermeable to both hydrophobic and large hydrophilic compounds (2). In addition to its role as a permeability barrier, β-barrel proteins and lipoproteins in the OM play key roles in a variety of other important processes, including motility, pathogenesis, and cell division (3). Because the periplasm lacks conventional sources of energy such as ATP, Gram-negative bacteria face a significant challenge in transporting and assembling OM components. To circumvent this challenge, cells utilize ATP hydrolysis in the inner membrane (IM) to transport LPS molecules across a protein bridge that spans the periplasm (4, 5). β-barrel proteins and lipoproteins also use ATP hydrolysis to cross the IM, but are escorted across the periplasm by soluble carriers (6, 7).

While relatively little is known about the transport of PLs to the OM, current understanding points to a mechanism that is highly distinct from the known OM transport pathways. Liposome fusion experiments in *Salmonella* Typhimurium demonstrated that unlike proteins and LPS, PL transport is bidirectional and indiscriminate (8). Rapid transfer from the OM to the IM was observed for all major and minor species of *Salmonella* PLs and even for cholesteryl oleate, which is not a normal component of bacterial membranes (8). One explanation consistent with these findings is that PLs can be transported by diffusional flow. Diffusive PL transport could occur at zones of hemifusion that form spontaneously. Diffusion could also require protein facilitators, for instance to encourage formation of hemifusions or to form protein channels through which PLs flow.

Although the bacterial PL transport pathway is currently unknown, the mechanisms by which cells maintain asymmetry in the OM are much better understood. When the integrity of the outer leaflet is disrupted, PLs from the inner leaflet migrate to fill gaps in the LPS, creating zones that are newly permeable to toxic hydrophobic compounds. The cell remedies this problem using the Mla (**M**aintenance of **l**ipid **a**symmetry) pathway, which removes mislocalized PLs from the outer leaflet and shuttles them to the IM (9). MlaA is a donut-shaped lipoprotein that sits in the outer membrane, removes PLs from the outer leaflet, and delivers them to the soluble carrier, MlaC. MlaC then transports them across the periplasm to the MlaFEDB complex, an ABC transport system that unloads MlaC and returns PLs to the IM.

In *E. coli*, a dominant negative mutation in *mlaA*, called *mlaA*\*, reverses the protein’s normal function (10). Instead of removing surface-exposed PLs, MlaA* allows properly localized PLs to flow through its pore into the outer leaflet (10, 11). Accumulation of PLs in the outer leaflet triggers a cell death pathway that results in lysis during stationary phase (10). First, the presence of PLs in the outer leaflet activates the OM phospholipase PldA, which cleaves surface-exposed PLs, generating breakdown products that signal to increase production of LPS (12). Hyper-production of LPS destabilizes the OM, resulting in loss of OM material through blebbing. PLs then flow from the IM to the OM to replace the lost material. In stationary phase, cells can no longer synthesize new PLs to replace those lost from the IM. As a result, PL flow causes the IM to shrink and ultimately rupture.

We hypothesized that changing the rate of PL flow from the IM to the OM would affect the rate of lysis in *mlaA*\* cells, since PL flow to the OM is what eventually causes the IM to rupture. Hence, we should be able to identify genes that affect PL transport through genetic interactions with *mlaA*\*. Our screen identified *yhdP*, a gene already known to play a role in maintaining the barrier function of the OM (13). Deletion of *yhdP* slowed lysis, but did not restore wild-type LPS levels, indicating that it affects a step in the pathway after LPS levels have already increased. Single-cell microscopy showed that the IM of *mlaA*\* Δ*yhdP* cells shrank more slowly, implying slower anterograde flow. In *mlaA*\* cells, PL flow ultimately leads to IM rupture but also compensates for loss of OM material while cells are actively growing. By contrast, without YhdP, the OM ruptured before the IM, suggesting that these cells cannot efficiently compensate for OM loss through anterograde flow.

## Results

### *mlaA*\* causes high-flux, passive phospholipid flow

It was previously shown that PL flow in *mlaA*\* cells is not affected by membrane depolarization or ATP synthase mutations, indicating that flow occurs via a passive mechanism (11). To further characterize this pathway, we quantified the rate of anterograde flow. We induced the cell division inhibitor SulA (14) and then transitioned exponentially growing cells onto agarose pads containing spent medium to cause the *mlaA*\* death phenotype. The SulA-induced cells became filamentous, and hence we could quantify the IM shrinkage from one of the poles prior to cell death (Figure 1A, white arrows) more easily than in non-filamentous cells. Since IM shrinkage in *mlaA*\* cells is the result of PL transport to the OM (10), we measured the rate of shrinkage as a proxy for the PL transport rate.

**Figure 1:**
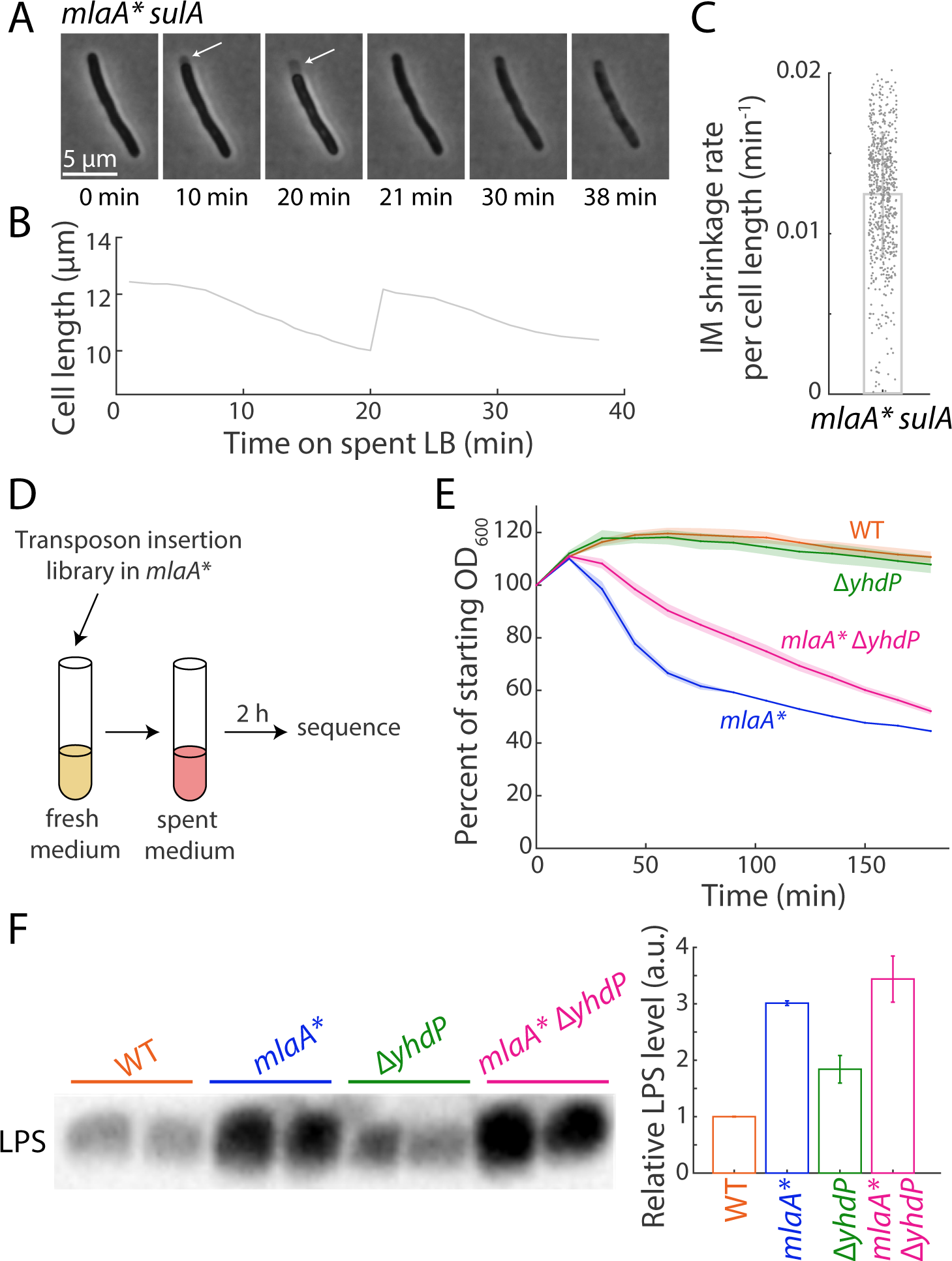
Deletion of *yhdP* slows the rate of *mlaA*\* lysis. A) Exponentially growing *mlaA*\* *sulA* cells were transitioned onto an agarose pad containing spent medium. In a typical cell, the IM shrank away from the cell wall (white arrows) before the cell eventually popped (*t*=21 min) and died. B) Cell length of the cell in (A) initially decreased, then rapidly snapped back to approximately the initial size at the time of transition to spent medium, and finally decreased due to leakage. C) During the initial 20 min in spent medium, the IM length shrank ~1% per minute. Each dot represents a single cell (total *n* = 677 cells), and the bar represents mean ± standard deviation (S.D.). D) Schematic of TraDIS selection. Libraries were grown into late exponential phase and transitioned to spent medium for 2 hours to induce lysis. The resulting library was subsequently sequenced for enrichment of mutants. E) Cultures were grown to late exponential phase (OD_600_~0.8), spun down, and resuspended in spent medium to induce lysis. OD_600_ was measured to determine rate of lysis. Deletion of *yhdP* slowed down *mlaA*\*-mediated lysis. Data points are mean ± S.D. with *n* = 3 replicates. F) Overnight cultures were normalized by OD_600_ and assayed for LPS abundance by immunoblotting. Left: immunoblotting gel image. Right: quantification of LPS abundances. Data points are mean ± S.D. with *n* = 2 biological replicates. Deletion of *yhdP* did not affect LPS levels either alone or in combination with *mlaA*\*.

The IM shrunk by ~20% in approximately 20 min (Figure 1B,C), corresponding to a PL flow rate of 1.2 ± 0.4% of the cell length per min. That a substantial fraction of the IM can be transported quickly even under the energetic limitations that occur upon entry into stationary phase provides further evidence that PL flow can occur via a diffusive mechanism. It also shows that the diffusive pathway is high-flux, permitting transport of a large proportion of the IM within a short period of time.

### Genetic interactions with *mlaA*\* depend on the length of time in spent medium

To identify genetic interactions with *mlaA*\*, we constructed transposon insertion libraries in *mlaA*\* and Δ*mlaA* cells. We grew the libraries to late exponential phase and incubated them in spent medium overnight to induce lysis. We repeated this process three times successively, inferring that the survival of any mutant that suppressed *mlaA*\*-mediated cell death would be amplified by the repeated incubations.

As expected, by far the most abundant hit was *mlaA*, since null mutations in *mlaA*\* prevent production of the mutant protein, completely suppressing cell death (10). The next most abundant hit was *pldA*, again expected as without PldA, there is no signal to increase production of LPS (10). After three rounds of incubation, insertions in *mlaA* and *pldA* accounted for 96.3% of all reads. Among the other hits (Table 1), several were known to affect LPS levels, corresponding to the results of a previous low-throughput screen, which also identified several suppressor mutations that lowered LPS levels (10). Since overproduction of LPS is a critical step in the cell-death pathway, mutations that restore wild-type LPS levels are expected to suppress lysis independent of any potential impact on PL transport (10). We therefore sought to find a genetic disruption that suppressed *mlaA*\* without lowering LPS levels.

**Table 1:** Percentage of reads in the *mlaA*\* library mapping to suppressor genes following three successive overnight incubations in spent medium.

| Gene name | % of total reads,<br>incubation 1 | % of total reads,<br>incubation 2 | % of total reads,<br>incubation 3 |
| --- | --- | --- | --- |
| *mlaA* | 48.1 | 84.1 | 86.0 |
| *pldA* | 14.7 | 9.8 | 10.3 |
| *lptC* | 8.6 | 0.9 | 0 |
| *dsbA* | 4.5 | 0 | 0 |
| *yaiP* | 1.3 | 2.7 | 1.7 |
| *acs* | 0.7 | 0 | 0 |
| *fadE* | 0.6 | 0 | 0 |
| *secA* | 0.5 | 0 | 0 |
| *asmA* | 0.3 | 0 | 0 |
| *yejM* | 0.2 | 0 | 0 |

Genomic DNA was extracted from the *mlaA*\* library following overnight incubation, which induces lysis, and transposon junctions were sequenced. Reads were mapped to the *E. coli* MC4100 genome and open reading frames were quantified to identify which gene disruptions were enriched after each incubation and hence were potential
suppressors. The two known strongest suppressors of *mldA*<sup>\*</sup>, *mldA* and *pldA*, quickly predominated in the culture.

**Table 2:** Number of reads in the *mlaA*\* and Δ*mlaA* libraries mapping to various genes after a two-hour incubation in spent medium.

| Gene name | # of reads – *mlaA*<sup>*</sup> | # of reads – $\Delta mlaA$ | $\log_2(mlaA^*/\Delta mlaA)$ |
| --- | --- | --- | --- |
| *yhdP* | 2,512,727 | 49,893 | 5.7 |
| *cyaA* | 1,153,018 | 83,061 | 3.8 |
| *mlaA* | 680,035 | N/A | N/A |
| *cysG* | 348,756 | 13,056 | 4.7 |
| *rbsD* | 215,556 | 13,903 | 4.0 |
| *sdhA* | 139,279 | 16,305 | 3.1 |
| *rssB* | 138,738 | 3885 | 5.2 |

Genomic DNA was extracted from the *mlaA*<sup>\*</sup> and $\Delta mlaA$ libraries following 2 h of incubation in spent medium, and transposon junctions were sequenced. Reads were mapped to the *E. coli* MC4100 genome and open reading frames were quantified. The most abundant gene disruption in the *mlaA*<sup>\*</sup> library was *yhdP*.

Since the most potent suppressors of *mlaA*\* block the earliest steps of the pathway, we hypothesized that slowing PL flow, the final step in the pathway, would only slow lysis. Hence, we carried out a similar experiment in which cells were only incubated for two hours in spent medium rather than overnight, to identify partial suppressors of *mlaA*\* (Figure 1D). Now, the most abundant hit in the *mlaA*\* library was *yhdP*, a large (1266 amino acid) IM protein. Interestingly, another member of its protein family, *asmA*, was identified as a suppressor in the previous screen (Table 1). YhdP has been shown to enhance OM permeability barrier function during stationary phase, but its mechanism is currently unknown (13). Deletion of *yhdP* causes sensitivity to SDS/EDTA and vancomycin regardless of growth phase, indicating that it plays a role in maintaining OM integrity (15). To confirm that disruption of *yhdP* inhibits lysis, we grew *mlaA*\* Δ*yhdP* cells to late exponential phase, resuspended them in spent medium, and measured OD over time; deletion of *yhdP* slowed the rate of lysis of *mlaA*\* cells (Figure 1E).

### Deletion of *yhdP* slows *mlaA*\* lysis without lowering LPS levels

Since modulating PL flow would affect a step in the cell death pathway after LPS levels have already increased, we expected that inhibiting PL flow would slow *mlaA*\* lysis without restoring wild-type LPS levels. To test the effects of *yhdP* deletion on *mlaA*\* cells, we measured LPS levels by immunoblotting (Methods). Deletion of *yhdP* had no effect on LPS levels either alone or in combination with *mlaA*\*, suggesting that it affects a later step in the pathway (Figure 1F). In addition, this finding suggests that *yhdP* does not slow lysis by affecting LPS transport, as it has been shown that slowing transport of LPS also reduces LPS levels (13).

### Deletion of *yhdP* slows shrinking of the IM

In *mlaA*\* cells, shrinking of the IM away from the cell pole is thought to reflect anterograde PL flow to the OM (10). We therefore expected that a mutation that slows PL flow would also slow IM shrinking. To determine whether *yhdP* deletion affects PL flow, we imaged *mlaA*\* or *mlaA*\* Δ*yhdP* cells during incubation in a microfluidic flow cell. Cells were first kept in LB until they reached steady-state growth, and then rapidly switched into spent medium. With continuous flow of spent medium, all *mlaA*\* cells died within 20-30 min (10). Deletion of *yhdP* delayed cell death (Figure 2A), consistent with the dynamics in bulk culture (Figure 1E).

**Figure 2:**
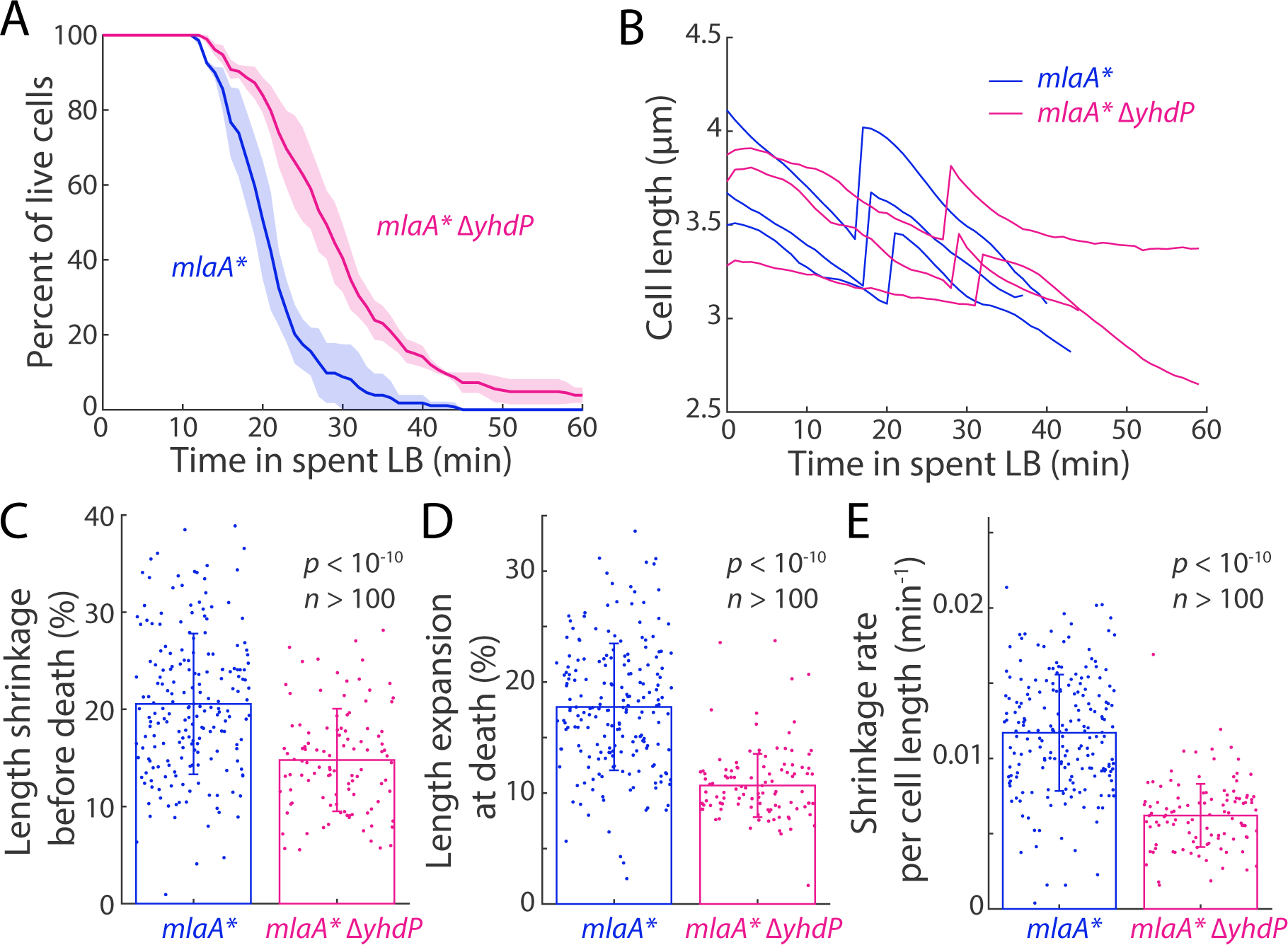
Deletion of *yhdP* slows shrinking of the IM during transition to spent medium. A) *mlaA*\* and *mlaA*\* Δ*yhdP* cells were separately incubated in a microfluidic flow cell and transitioned from fresh LB to spent medium to induce cell death. Consistent with bulk measurements, deletion of *yhdP* slowed down cell death. Data points are mean ± S.D. with *n* = 3 replicates of at least 50 cells in each experiment. B) Representative single-cell traces after switching to spent medium. C) Deletion of *yhdP* reduced total shrinkage in *mlaA*\* cells by ~50% (*p* < 10^10^, *n* > 100 cells, two-tailed Student’s *t*-test). D) During the “popping” immediately preceding lysis, *mlaA*\* and *mlaA*\* Δ*yhdP* cells returned to approximately their initial length prior to the transition to spent medium (compare length expansion to the shrinkage in (C); *mlaA*\* cells exhibited more expansion than *mlaA*\* Δ*yhdP* cells, *p* < 10^10^, *n* > 100 cells, two-tailed Student’s *t*-test). E) Deletion of *yhdP* slowed down the shrinkage rate of *mlaA*\* cells (*p* < 10^10^, *n* > 100 cells, two-tailed Student’s *t*-test). In (C-E), each dot represents a single cell, and the bar plots represent mean ± S.D.

For both strains, the cytoplasm started to shrink immediately upon the transition to spent medium. The cytoplasm of *mlaA*\* cells shrank by an average of ~0.7 μm (20%, Figure 2B,C) over ~20 min (Figure 2A) and appeared to increase in density, followed by a “popping” expansion and then gradual loss of phase contrast (Figure 2B) that we previously characterized as typical of *mlaA*\*-mediated death (10). *mlaA*\* Δ*yhdP* cells displayed a qualitatively similar death trajectory (Figure 2B). The average time to lysis was longer (29 min, Figure 2A) and yet less shrinkage occurred (0.5 μm, 15%, Figure 2B,C) before popping than in *mlaA*\* cells. In both strains, the expansion at cell death roughly restored cell length to the pre-shrinkage size (Figure 2C,D), suggesting that the cell envelope returned to a relaxed state after the expansion. Shrinkage rate prior to popping was also slowed down in *mlaA*\* Δ*yhdP* cells by 50% (Figure 2E). Taken together, these data indicate that YhdP plays an important role in PL transport during *mlaA*\*-mediated lysis.

### The effect of YhdP on lysis is cyclic ECA-independent

It was previously shown that the OM permeability phenotypes of Δ*yhdP* cells can be suppressed by preventing synthesis of cyclic enterobacterial common antigen (ECA), indicating that YhdP regulates cyclic ECA (15). To test whether the effect of *yhdP* deletion on *mlaA*\*-mediated lysis also depends on cyclic ECA, we constructed strains lacking *wzzE*. WzzE is the ECA chain length regulator, and in its absence cyclic ECA is not synthesized. If the effect of *yhdP* deletion on lysis rate also depends on cyclic ECA, we would expect that deleting *wzzE* in *mlaA*\* Δ*yhdP* cells would reverse the effect of *yhdP* deletion, resulting in dynamics upon transition to spent medium similar to that of *mlaA*\* alone.

To quantify the effect of cyclic ECA in *mlaA*\* cells, we imaged *mlaA*\* Δ*yhdP* Δ*wzzE* cells in a microfluidic device during the transition to spent medium. Deletion of *wzzE* did not restore *mlaA*\*-like death dynamics (Figure 3A), nor did it change the shrinkage rate of the *mlaA*\* Δ*yhdP* strain (Figure 3B). Deletion of *wzzE* did not affect the death (Figure 3A) or shrinkage (Figure 3B) of *mlaA*\* cells, indicating that the effect of YhdP on PL transport during *mlaA*\*-mediated lysis does not require cyclic ECA.

**Figure 3:**
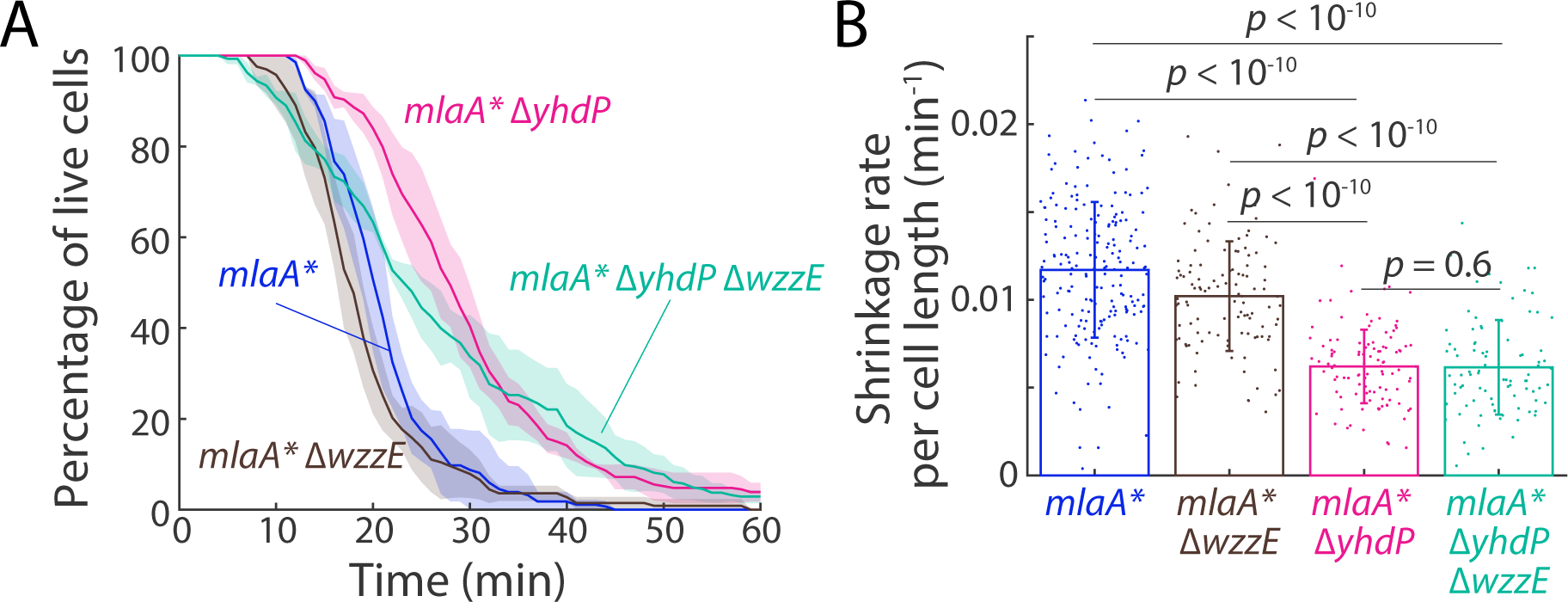
Cyclic ECA is not responsible for suppression of death by Δ*yhdP*. A) Cells were incubated in a microfluidic flow cell and transitioned from fresh LB to spent medium to induce cell death. Deletion of the ECA biosynthesis gene *wzzE* did not restore *mlaA*\*-like lysis dynamics to *mlaA*\* Δ*yhdP. mlaA*\* Δ*yhdP* Δ*wzzE* cells exhibited distinct and slower death dynamics compared to *mlaA*\* cells, while deletion of *wzzE* from *mlaA*\* slightly accelerated cell death. Data points are mean ± S.D. with *n* = 3 replicates. B) Deletion of *wzzE* did not alter the shrinkage rate of *mlaA*\* Δ*yhdP* cells (*n* > 100 cells) and only slightly reduced the rate in *mlaA*\* cells, indicating that the effect of YhdP on lysis is cyclic ECA-independent. Each dot represents a single cell (*n* > 100 cells for each strain), and the bar plots represent mean ± S.D. *p*-values are from two-tailed Student’s *t*-tests.

### Deletion of *yhdP* weakens the OM chemically and mechanically

Another explanation for how deletion of *yhdP* could slow lysis is by preventing loss of OM material. To test whether deleting *yhdP* improves OM integrity in *mlaA*\* cells, we assayed OM permeability by plating on vancomycin or SDS/EDTA. It was previously shown that cells lacking *yhdP* are vancomycin-sensitive (15). However, by plating on a low concentration of vancomycin such that wild-type, *mlaA*\*, and Δ*yhdP* cells all grew to the same dilution as on LB without drug, we observed that *mlaA*\* Δ*yhdP* cells had a synthetic OM permeability defect (Figure 4A). On SDS/EDTA, *mlaA*\* and Δ*yhdP* were both sensitive; combining the two mutations did not relieve the defect (Figure 4A). These results demonstrate that deletion of *yhdP* does not slow lysis by enhancing OM integrity.

**Figure 4:**
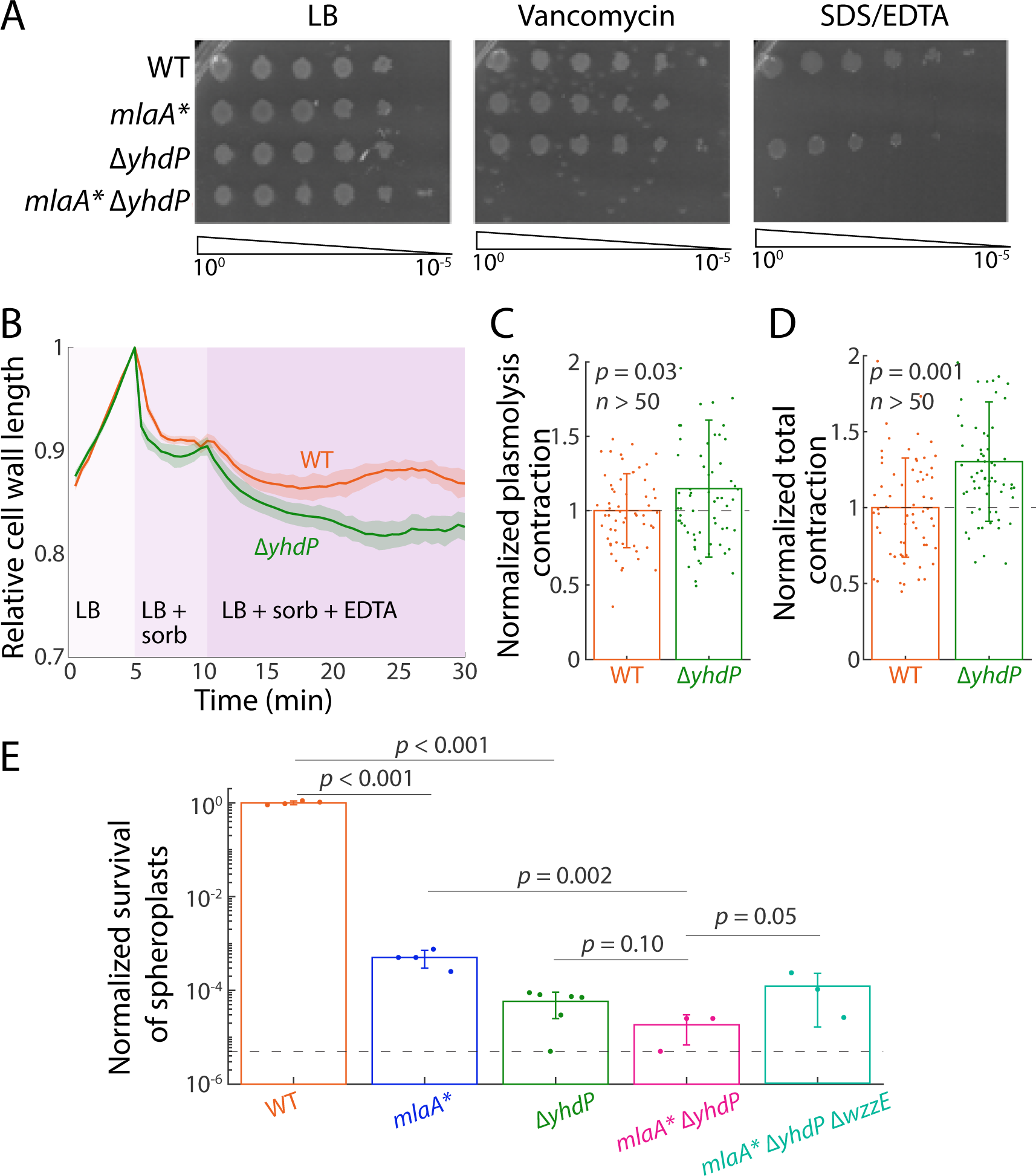
Deletion of *yhdP* chemically and mechanically disrupts the OM. A) Overnight cultures were normalized by OD, serially diluted, and plated on LB, LB + 20 μg/mL vancomycin, and LB + 0.5% SDS/0.5 mM EDTA. *mlaA*\* and Δ*yhdP* have a synthetic permeability defect with vancomycin, and neither *mlaA*\* nor *mlaA*\* Δ*yhdP* cells grew with SDS/EDTA. B) Exponentially growing cells were loaded into a microfluidic device and allowed to grow in LB before being exposed to a large hyperosmotic shock with 3 M sorbitol, then treated with EDTA in the precence of sorbitol. The length of the fluorescently labeled cell wall was tracked. Sorbitol treatment relieved turgor pressure and reduced cell-wall length. EDTA treatment disrupted the OM and led to a further decrease in cell length. In both conditions, Δ*yhdP* cells shrank more compared to wild-type cells. C,D) Length contraction upon sorbitol (C) and EDTA (D) treatment for cells in (B). In both conditions, Δ*yhdP* cells shrank more than wildtype, indicating a mechanically weakened OM. Individual dots are data from single cells (*n*>50 for each strain), and bar plots represent mean ± S.D. *p* values are from a two-tailed Student’s *t*-test. E) Spheroplasts were generated overnight in the presence of cefsulodin, and then washed and plated on fresh media. Both *mlaA*\* and Δ*yhdP* exhibited a mechanically weakened OM and reduced spheroplast survival rates. Deletion of *wzzE* partially rescued the mechanical defect in *mlaA*\* Δ*yhdP* cells. Dots represent biological replicates (*n*>3 replicates for each strain), and the bar plots are mean ± S.D. *p* values are from one-tailed Student’s *t*-test.

Since deleting *yhdP* increased OM permeability in *mlaA*\* cells, we wondered whether inhibition of anterograde flow might be due to destabilization of the OM. To further characterize the effect of YhdP on the OM, we investigated its impact on OM mechanical strength. In a previous study, we showed that the mechanical stiffness of the *E. coli* OM is greater than or comparable to that of the cell wall, and that genetic or chemical perturbations to the OM can reduce the overall stiffness of cells (16). To determine if YhdP plays a role in determining OM stiffness, we utilized an assay in which exponentially growing cells are first exposed to a large, hyperosmotic shock with 3 M sorbitol, and then treated with EDTA. We used a microfluidic flow cell to precisely control the timing of treatments and track single cells throughout (Methods). Upon the shock, wild-type cells experienced a large decrease in the length of the fluorescently labeled cell wall (Figure 4B), as expected since turgor pressure was relieved and hence the cell wall-OM envelope complex was no longer under stress. EDTA treatment, which disrupts the OM by rapidly inducing loss of LPS molecules (17, 18), led to a further decrease in cell length (Figure 4B), signifying that the stiff OM was holding the cell wall out beyond its rest length before its removal. Application of this assay to Δ*yhdP* cells showed greater contraction of the cell wall after the osmotic shock (Figure 4B,C) and after EDTA treatment (Figure 4B,D), indicating that the overall stiffness of Δ*yhdP* cells was lower than that of wildtype.

To further test whether deletion of *yhdP* weakened cells mechanically, we quantified the yield of viable cells after breaking down the cell wall using beta-lactam antibiotics to form wall-less spheroplasts with intact IM and OM (Methods). We previously showed that spheroplast yield is strongly correlated with the stiffness of the OM across chemical and genetic perturbations (16). In this assay, spheroplasts were generated overnight in the presence of cefsulodin, and then were washed and plated on fresh medium without antibiotics after the cell wall was removed. Survival in the absence of a cell wall relies on having a stiff outer membrane to bear the stress of turgor. We observed that *mlaA*\* and *yhdP* deletion each caused a dramatic (>1000-fold) decrease in spheroplast viability in comparison with wildtype (Figure 4E). The *mlaA*\* Δ*yhdP* double mutant exhibited a further decrease in spheroplast viability, highlighting the importance of YhdP in determining OM stiffness. However, deletion of *wzzE* partially suppressed the decrease in spheroplast viability due to Δ*yhdP* (Figure 4E), demonstrating that the effect of YhdP on OM mechanical strength is cyclic ECA-dependent.

Taken together, these results suggest that deleting *yhdP* does not slow lysis by preventing loss of OM material. Deletion of *yhdP* severely disrupts OM integrity, which is more likely to promote loss of OM material than to prevent it. Furthermore, *yhdP* deletion still slows lysis even when its effect on the mechanical strength of the OM is suppressed (Figure 3A,B, Figure S1), indicating that *yhdP’s* effect on lysis is not a result of its effect on OM mechanics.

### Impairment of phospholipid flow leads to OM rupture

We observed that the IM of *mlaA*\* Δ*yhdP* cells shrank more slowly and less relative to *mlaA*\* (Figure 2B-E). We would expect that a mutation that decreases PL flow would cause the IM to shrink more slowly. To explain why the IM shrank less before lysis, we wondered whether, in these cells, lysis occurs for a reason other than IM rupture. In *mlaA*\*, anterograde flow leads to rupture of the IM, followed shortly by OM rupture (10). We surmised that impairing PL flow in *mlaA*\* cells would increase the stress on the OM, potentially causing the OM to rupture before the IM.

To test this hypothesis, we constructed *mlaA*\* and *mlaA*\* Δ*yhdP* strains expressing both a cytoplasmic and a periplasmic fluorescent protein. When the *mlaA*\* strain was shifted into spent medium, shrinkage of the IM led to a large periplasmic space with high mCherry signal (Figure 5A, white arrow). The mCherry signal remained intact throughout shrinkage, and when the cells popped and lysed, periplasmic mCherry and cytoplasmic GFP signals were lost simultaneously in every cell (Figure 5A,B), presumably because rupture of the IM also led to rapid OM rupture (10). By contrast, in *mlaA*\* Δ*yhdP* cells, the extent of IM shrinkage was much smaller (Figure 2C), and the periplasmic mCherry signal remained largely uniform around cell periphery rather than intensified at a cell pole(s) (Figure 5C). During the transition to spent medium, the mCherry signal was lost tens of minutes before popping (Figure 5C,D), while the cytoplasmic YFP signal remained intact until popping occurred (Figure 5C). Taken together, these data indicate that disruption of anterograde flow caused by *yhdP* deletion in *mlaA*\* Δ*yhdP* cells leads to rupture of the OM before the IM (Figure 5E).

**Figure 5:**
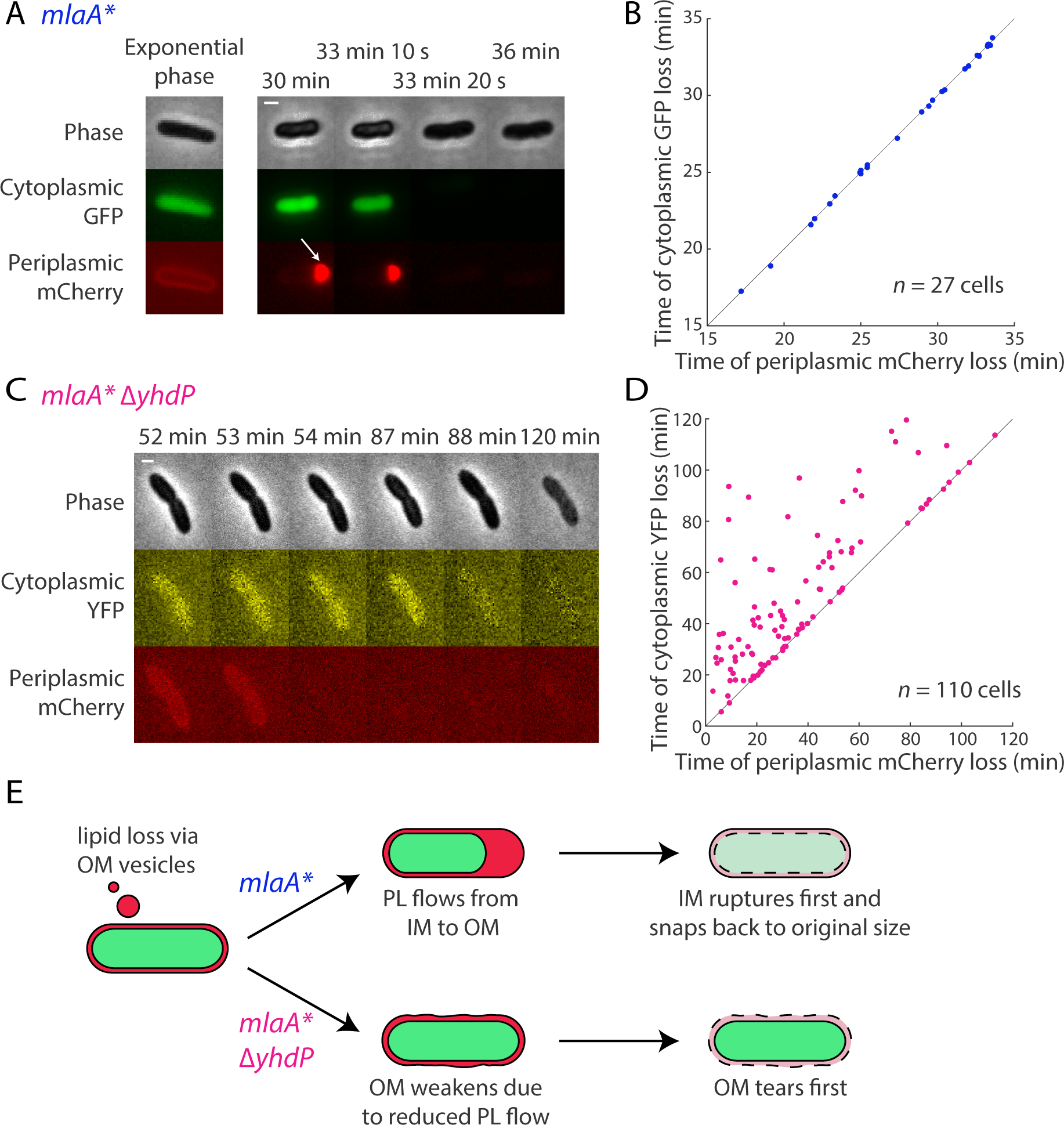
Deletion of *yhdP* from *mlaA*\* cells causes the OM to rupture before the IM in spent medium. A) Death trajectory of *mlaA*\* cells on an agarose pad with spent medium. Cells were labelled with periplasmic mCherry and cytoplasmic GFP. During shrinkage, PLs flowed from the IM to the OM, causing the IM to shrink away from the cell wall and OM. As a result, periplasmic mCherry was enriched at one cell pole (white arrow). At the time of “popping,” both fluorescence signals were lost in the same frame. Scale bar is 1 μm. B) During the transition to spent medium, mCherry and GFP signals were lost simultaneously in all *mlaA*\* cells (*n*=27). Dots represent single cells, and black line is *x = y*. The dots are slightly jittered to visualize overlapping data. C) Death trajectory of *mlaA*\* Δ*yhdP* cells on an agarose pad with spent medium. Cells were labelled with periplasmic mCherry and cytoplasmic YFP. During the period of shrinkage (52-87 min), the IM did not shrink away from cell wall and OM, as shown by the uniform mCherry signal around the cell periphery. The cell also lost its periplasmic mCherry signal tens of minutes before losing cytoplasmic YFP signal, suggesting that the OM ruptured before the IM. Scale bar is 1 μm. D) During the transition to spent medium, the mCherry signal was lost at least 2 min before the YFP signal in *n*=74 (out of 110) *mlaA*\* Δ*yhdP* cells, indicating that deletion of *yhdP* leads to rupture of the OM before the IM. In all other cells both signals were lost simultaneously. Dots represent single cells, and black line is x = *y*. The dots are slightly jittered to visualize overlapping data. E) Model of Δ*yhdP*-mediated death. The *mlaA*\* mutation leads to membrane loss via OM vesicles and disrupts PL homeostasis during the transition into stationary phase. In the *mlaA*\* background (top), PLs flow from the IM to the OM to replenish the membrane loss, causing the IM to shrink away from OM, and eventually leading to cell death through IM rupture. By contrast, in *mlaA*\* Δ*yhdP* cells (bottom), deletion of *yhdP* suppresses PL flow, leading to further weakening of an already compromised OM that ruptures before the IM.

## Discussion

The existence of fusion junctions facilitating PL flow between the IM and OM has been a matter of controversy for some time. In the 1960s, electron microscopy showed sites of contact between the two membranes, but improved microscopy methods called into question the existence of these “Bayer’s junctions” (19, 20). While it may be the case that the junctions observed in those early images were indeed artifacts, several lines of evidence now suggest that intermembrane PL transport can occur via diffusion.

Previous studies showed that PL transport is bidirectional and can involve even non-native lipids (8, 21). In the *mlaA*\* mutant, PL flow does not require either ATP or proton motive force (11). In addition, in this mutant approximately 20% of the IM is lost by transport even under the nutrient limitations that trigger entry into stationary phase. These data are strong evidence that PL flow in *mlaA*\* cells is passive, and occurs through a high-flux pathway. It remains to be seen whether this pathway functions in normal PL transport or is active only in certain conditions.

In this study, we provide evidence that YhdP is involved in modulating the high-flux PL transport pathway. Time-lapse imaging showed that deleting *yhdP* slowed shrinkage of the IM in *mlaA*\* cells (Figure 2), implying that PLs flowed more slowly from the IM to the OM. In *mlaA*\*, loss of lipids from the IM ultimately causes it to rupture (10). As a result, slowing PL flow delays cell death. However, since PL flow also compensates for loss of OM material, slowing flow from the IM comes at the cost of OM integrity. Thus, while lysis takes longer in *mlaA*\* Δ*yhdP* cells, when it does occur, the OM rather than the IM ruptures first (Figure 5D).

Cells survive without *yhdP*, suggesting that YhdP functions specifically in high-flux PL transport. If it does play a role in normal PL transport, then there must be multiple, redundant pathways. How YhdP modulates PL transport is still unknown, but an intriguing possibility is suggested by its protein family. YhdP belongs to a family of six “AsmA-like” proteins (AsmA, TamB, YdbH, YicH, YhjG, and YhdP). Two members of this family, AsmA and TamB, are predicted to share homology with the eukaryotic PL transporter, Vps13 (22). Vps13 forms a hydrophobic channel through which PLs are transported between membranes (23, 24). The structure of TamB also includes a channel with a highly hydrophobic interior (25). Interestingly, it has been suggested that due to its ability to accommodate many lipids at once, Vps13 functions specifically in high-flux PL transport (26).

While our study does not determine YhdP’s molecular mechanism, it does rule out certain possibilities. Deleting *yhdP* does not lower LPS levels (Figure 1F), hence it must affect a step in the *mlaA*\* death pathway after LPS levels have already increased. Moreover, the effect of *ydhP* deletion on *mlaA*\* lysis cannot be explained by slowed transport of LPS to the OM, as it has previously been shown that slowing LPS transport also decreases LPS levels (12). It is also unlikely that deleting *yhdP* slowed lysis (Figure 2A,B) by preventing loss of OM material, as *yhdP* deletion has a severe negative impact on OM integrity (Figure 4). Of the remaining options, a direct role in transport is certainly the simplest. YhdP is a large (1266 amino acid) IM protein with one clear N-terminal and possibly a second, C-terminal transmembrane domain. Given the size of its periplasmic domain, it is plausible that YhdP can span the periplasm, but further structural and biochemical studies are needed to determine its precise role in anterograde PL transport. Regardless, our data provide new insight into the process of PL flow and cell lysis caused by the dominant negative *mlaA*\* allele, and shed light on the multiple roles played by YhdP in the maintanence of OM integrity. The fact that YdhP changes both OM stiffness and permeability suggests an intriguing link between these two properties. Our discovery of a mutant capable of slowing PL transport should provide a useful foothold in the investigation of this poorly understood pathway.

## Methods

### Bacterial strains

The strains used in this study are listed in Table S1. Strains were constructed by generalized P1 transduction with all deletions originating from the Keio collection (27, 28). Kanamycin resistance cassettes were removed using the Flp recombinase system, as previously described (29). Overnight cultures were grown at 37 °C in lysogeny broth (LB) medium supplemented with 10 mM MgSO_4_ to prevent *mlaA*\* lysis and diluted into unsupplemented LB for subsequent experiments. When necessary, media were supplemented with 25 μg/mL kanamycin or 25 μg/mL tetracycline.

### TraDIS sample preparation

Transposon mutant libraries were constructed using the EZ-Tn5<KAN-2>TnP Transposome Kit (Epicentre) according to the manufacturer’s instructions. When preparing electrocompetent cells, overnight cultures were grown in LB supplemented with 5 mM MgSO_4_ to prevent lysis of *mlaA*\* and then subcultured in 2xYT medium. Following electroporation, cells were plated on LB+25 μg/mL kanamycin plates supplemented with 5 mM MgSO_4_. Approximately 300,000 and 150,000 colonies were pooled to construct the *mlaA*\* and Δ*mlaA* libraries, respectively. Genomic DNA was extracted from samples of 2×10^9^ cells after lysis using the DNeasy Blood and Tissue Kit (Qiagen) according to the manufacturer’s instructions. Libraries were prepared according to the TraDIS method (30) and sequenced on Illumina HiSeq 2500 Rapid flowcells as single-end, 75-nucleotide reads.

### TraDIS data analysis

Sequencing reads were mapped to the *E. coli* K12 genome using BWA v. 1.2.3. Mapped reads were quantified using htseq-count v. 0.6.0. The Integrative Genomics Viewer was used to visualize the mapped reads.

### Lysis curves

To generate spent medium, wild-type (MC4100) cultures were grown for 24 h in LB at 37 °C, cells were pelleted, and the supernatant was filter-sterilized using a 0.2-μm filter. All experiments were conducted using wild-type spent medium. To assay the rate of lysis, cultures were grown until OD_600_~0.8, pelleted, and resuspended in spent medium. Cultures were then incubated at 37 °C and OD_600_ was measured at 15-min intervals.

### Immunoblot analyses

The equivalent of 1 mL of culture at OD_600_~1 was taken from overnight cultures, pelleted, and resuspended in LDS sample buffer (Invitrogen). Samples were boiled for 10 min and allowed to cool. Samples were loaded on 4-12% SDS/polyacrylamide gel electrophoresis (PAGE) gels and run at 100 V. LPS was then transferred to nitrocellulose membranes and blocked in 5% non-fat dried milk for 1 h at room temperature. Membranes were then incubated overnight at 4 °C with anti-LPS antibody (1:400,000; Hycult Biotech) in milk. Membranes were washed and incubated with secondary antibody for 1 h at room temperature (1:20,000; Goat Anti-Mouse IgG (H+L)-HRP Conjugate; Bio-Rad).

### Efficiency of plating assay

Cultures were grown overnight in LB + 10 mM MgSO_4_, standardized by OD, and serially diluted. Dilutions were then transferred to plates using a 96-well-plate replica plater and incubated overnight at 37 °C.

### Single-cell imaging

Cells were imaged on a Nikon Eclipse Ti-E inverted fluorescence microscope with a 100X (NA 1.40) oil-immersion objective (Nikon Instruments). Images were collected on a DU885 electron-multiplying charged couple device camera (Andor Technology) or a Neo sCMOS camera (Andor Technology) using μManager version 1.4 (http://www.micro-manager.org) (31). Cells were maintained at 37 °C during imaging with an active-control environmental chamber (HaisonTech).

For experiments conducted on agarose pads, 1 μL of cells was spotted onto a pad of 1% agarose in fresh LB or spent medium. For transition experiments, exponentially growing cells were washed three times in spent medium before spotting. Flow-cell experiments were performed in ONIX B04A microfluidic chips (CellASIC) and medium was exchanged using the ONIX microfluidic platform (CellASIC).

### Imaging in microfluidic devices

Overnight cultures were diluted 100-fold into 1 mL of fresh LB and incubated for 2 h with shaking at 37 °C. B04A plates were loaded with medium and pre-warmed to 37 °C. Cells were loaded into the plate, which was incubated at 37 °C, without shaking for 30 min before imaging. As necessary, the cell envelope was stained with wheat germ agglutinin-AlexaFluor488 (WGA-AF488, Life Technologies), which was added to the loading well to a final concentration of 10 μg/mL prior to loading cells into the imaging chamber. The osmolarity of the growth medium was modulated with sorbitol (Sigma).

During plasmolysis/lysis experiments to quantify the effect of *yhdP* deletion on cell stiffness, cells were allowed to grow for 5 min in medium in the imaging chamber before being plasmolyzed with LB + 3 M sorbitol and exposed to LB + 3 M sorbitol + 10 mM EDTA 5 min later.

### Image analysis

Time-lapse images were first segmented with the software *DeepCell* (32), and the resulting segmented images were analyzed using *Morphometrics* (33) to obtain cell contours at sub-pixel resolution. Static images were directly segmented using *Morphometrics* (33). Cell width and length were calculated using the MicrobeTracker meshing algorithm (34).

### Quantification of spheroplast viability and growth

Overnight cultures of the appropriate strains were diluted 1:100 into LFLB (LB supplemented with 3.6% sucrose and 10 mM MgSO_4_). Cultures were incubated at 37 °C for 1 h, normalized to OD_600_ ~ 0.08, at which point cefsulodin was added to a final concentration of 60 μg/mL. Cells were further incubated for 12 h with shaking at 30 °C. Ten microliters of serial ten-fold dilutions were plated on LFLB plates. Plates were incubated at 30 °C for 24 h, and colony-forming units were counted manually.

## Acknowledgements

The authors thank the Huang and Silhavy labs for helpful suggestions and the Genomics Core Facility of Princeton University for next-generation sequencing experiments. This research was supported by the National Institute of General Medical Sciences of the National Institutes of Health under grants 5R35GM118024 (to T.J.S.) and T32-GM007388 (to J.G.). The authors acknowledge partial support from NIH Grant R01 GM082938 (N.S.W.) and support from a James McDonnell Postdoctoral Fellowship (to H.S.). K.C.H. is a Chan Zuckerberg Biohub Investigator.

## Supplementary Tables

**Table S1:** Strains used in this study.

| *Escherichia coli* K-12 strains | Genotype and relevant features | Reference |
| --- | --- | --- |
| MC4100 | F-*araD139 (argF-lac)*U169<br>*rpsL150 relA1 flb5301 deoC1*<br>*ptsF25 thi* | (35) |
| HC735 | MC4100 ara+ $\Delta yfdI$ | (10, 36) |
| HC687 | MC4100 ara+ $\Delta yfdI mlaA^*$ | (10, 27) |
| HC736 | MC4100 ara+ $\Delta yfdI \Delta mlaA$ | (10, 27, 37) |
| JG153 | MC4100 ara+ $\Delta yfdI$<br>$\Delta yhdP::kan$ | (10, 27) and this study |
| JG152 | MC4100 ara+ $\Delta yfdI mlaA^*$<br>$\Delta yhdP::kan$ | (10, 27) and this study |
| JG174 | MC4100 ara+ $\Delta yfdI$<br>*intS::tn10* | This study |
| JG178 | MC4100 ara+ $\Delta yfdI mlaA^*$<br>*intS::tn10* | This study |
| JG176 | MC4100 ara+ $\Delta yfdI$ *mlaA* *<br>*intS::tn10* $\Delta yhdP$ | (27, 37) and this study |
| JG179 | MC4100 ara+ $\Delta yfdI$ *mlaA* *<br>*intS::tn10* $\Delta yhdP$<br>$\Delta wzzE::kan$ | (27, 37) and this study |
| JG197 | MC4100 ara+ $\Delta yfdI$ *mlaA* *<br>*intS::tn10* $\Delta wzzE::kan$ | (27, 37) and this study |
| KC637 | HC687 pMT37-*sulA* | (10) |
| KC626 | HC687 attHK:*Plac-mCherry*<br>pZS21-*GFP* | (10) |
| KC1307 | JG153 attHK:*Plac-mCherry*<br>*Plac-YFP* | (10) and this study |

## Notes

### Competing Interest Statement

The authors have declared no competing interest.

